# miRstring: An RNA language model enables mature miRNA decoding and artificial small RNA design across species

**DOI:** 10.64898/2026.09.17.752258

**Authors:** Ruikang Peng, Xiang Li, Yitian Fang, Xiang Yu

**Affiliations:** School of Agriculture and Biology, Shanghai Jiao Tong University, Shanghai 200240, China; School of Life Sciences and Biotechnology, Shanghai Jiao Tong University, Shanghai 200240, China; School of Pharmaceutical Sciences, Shanghai Jiao Tong University, Shanghai 200240, China

**Keywords:** RNA language models, microRNAs, artificial miRNA design, artificial small RNAs

## Abstract

**Introduction:** MicroRNAs (miRNAs) are processed from structured precursors and subsequently loaded into Argonaute proteins to repress target mRNAs. Accurate decoding of mature miRNAs from precursor sequences is fundamental to miRNA annotation and the rational engineering of artificial miRNAs. However, existing computational approaches are largely developed for humans, with limited ability to generalize across diverse plants, animals, and viruses. Moreover, a unified data-driven framework that links mature-miRNA decoding with the predictive design of artificial miRNA precursors is still lacking, restricting the scalable application of miRNA biotechnology across species.

**Objectives:** We develop and evaluate miRstring, a biogenesis-aware RNA language framework for decoding the four boundaries of mature miRNAs from precursor context across species, and further investigate its utility for cross-species miRNA annotation and rational artificial miRNA design.

**Methods:** miRstring, a biogenesis-aware RNA language framework, was developed using 77,708 miRNA precursors spanning 414 species. The model was evaluated against existing methods under both family-held-out and species-held-out settings to assess its ability to generalize to unseen miRNA families and species. Attention-based analyses were further performed to examine whether miRstring captured biologically relevant features associated with miRNA processing, and the framework was subsequently applied to predictive design of artificial miRNA precursors.

**Results:** miRstring substantially outperformed existing methods across species-held-out evaluations and, notably, retained strong performance under the more stringent family-held-out setting. It accurately identified mature-miRNA start sites, while attention was enriched at endonuclease-cleaved miRNA/miRNA* boundaries, revealing biologically meaningful processing features. Importantly, miRstring-guided precursor design markedly enhanced artificial miRNA-mediated target repression, demonstrating its utility for rational miRNA engineering.

**Conclusion:** miRstring provides a scalable framework linking cross-species mature-miRNA annotation with predictive artificial miRNA design, extending computational miRNA analysis toward artificial intelligence-driven small-RNA biotechnology.

## 1. Introduction

MicroRNAs (miRNAs) are endogenous small noncoding ribonucleic acids (RNAs) that post-transcriptionally regulate complementary mRNAs by repressing translation and promoting mRNA decay[1,2]. As mature guide RNAs loaded into Argonaute (AGO) proteins, miRNAs direct the miRNA-induced silencing complex (miRISC) to target transcripts through sequence complementarity, thereby reducing gene output through translational repression and mRNA destabilization; in plants, strict guide–target complementarity can additionally lead to AGO-mediated target cleavage[2–5]. Because target recognition is specified by the mature miRNA sequence itself, the precise identity of the mature product determines which transcripts, pathways and regulatory programs can be engaged[3,6,7]. Accurate precursor-to-mature definition is therefore a prerequisite for downstream tasks such as miRNA catalogue expansion, expression quantification, cross-species comparison and target inference[7,8]. Yet recovering mature miRNA sequences from precursors remains laborious, typically requiring multi-step small RNA-seq analysis, while cleavage-site–level confirmation often depends on additional experimental validation such as rapid amplification of cDNA ends (RACE), cross-linking and immunoprecipitation (CLIP)-based assays and perturbation of core biogenesis factors[5,8–11]. As a result, rapid and reliable precursor-to-mature prediction remains both difficult and necessary[7,8].

miRNA maturation proceeds through related but not identical intracellular routes across different biological groups[4,7]. In animals, a primary microRNA (pri-miRNA) hairpin is first processed in the nucleus by the ribonuclease III (RNase III) Microprocessor complex, Drosha–DGCR8, producing a pre-miRNA that is exported to the cytoplasm and further cleaved by Dicer into an approximately 22-nt duplex[7,12]. In plants, analogous processing takes place largely in the nucleus through Dicer-like 1 (DCL1) together with HYL1 and SERRATE, so that both cleavage steps are completed without a separate cytoplasmic stage[4]. Several viruses also encode miRNAs and can access host or non-canonical processing pathways, adding further diversity to the precursor-to-mature transition[13–15]. Across these systems, processing yields a miRNA duplex, typically comprising a mature guide miRNA which is loaded into Argonaute and mediates silencing and a passenger strand (historically denoted miRNA*) which is usually displaced and degraded after loading, with characteristic overhangs. Guide-strand selection is then influenced by duplex end stability and 5′ nucleotide identity[11,16–18]. Defining a mature miRNA from its precursor therefore requires resolving the cleavage positions that delimit the final duplex along a structured hairpin and the mature sequence ultimately presented to Argonaute[7,12].

Despite progress in miRNA cleavage-site prediction, existing models—from cleavage-pattern classifiers built with support vector machines (SVMs) or gradient boosting (PHDcleav, LBSizeCleav, ReCGBM) and recurrent neural network (RNN)-based predictors such as DeepMirCut to more recent deep-learning models such as DiCleave and DiCleavePlus trained on relatively small, human-centric benchmarks—have primarily been developed and evaluated under in-distribution settings with limited precision and thus do not establish robust generalization across unseen species and miRNA families[19–23].

To fill this gap, we introduce miRstring, a multi-module deep-learning model that predicts mature miRNA duplexes directly from precursor sequences. We train and evaluate miRstring on a curated dataset spanning 414 species, 77,708 precursor records and 16,654 miRNA families, using strict out-of-distribution benchmarks that hold out entire species and families. With data augmentation, miRstring not only ingests both sequence and secondary-structure features, but integrates them via a structure-guided cross-attention module, and predicts the precise start and end boundaries of both arms of the mature miRNA duplex. In parallel, a lightweight binary head attached to the shared representation provides a precursor-validity estimate, while a downstream assignment module determines which recovered arm is the guide strand and which is the passenger strand. Moreover, miRstring offers interpretability of miRNA predictions derived from attention-guided sequence– structure insights, enabling post-hoc analyses of the local sequence and pairing features associated with predicted duplex boundaries. A low-dimensional embedding of final-layer head features further separates animal from plant precursors, consistent with lineage-specific processing context.

Building on miRstring’s recovery of mature miRNA sequences from precursor context, we also developed a model-guided approach for artificial miRNA (amiRNA) design. Current amiRNA design often relies on reusing a small set of endogenous precursor scaffolds while replacing only the miRNA duplex, but this scaffold-reuse strategy poses two persistent problems: processing efficiency depends strongly on the compatibility between scaffold and designed duplex, and cleavage imprecision can alter the recovered mature sequence, especially at the 5′ end, thereby affecting guide identity and increasing off-target risk[24–27]. By capturing boundary-associated sequence–structure signals, miRstring enables a scaffold-aware, model-guided route for amiRNA design in Metazoa and Viridiplantae, prioritizing candidate backbones for user-specified mature sequences while preserving native precursor context as much as possible.

## 2. Material and Methods

### Data collection and evaluation design

miRNA precursor annotations were integrated from miRBase, MirGeneDB and PmiREN. Records were retained when an annotated mature product was available and precursor length did not exceed 250 nt, yielding 77,708 precursors spanning 414 species. For curated clade-specific evaluation, 21 metazoan species present in MirGeneDB but absent from miRBase and 10 plant species present in PmiREN but absent from miRBase were reserved as unseen-species sets. Family-held-out datasets were generated by assigning all members of the same miRNA family to a single split using family-size-stratified partitioning, thereby preventing family-level leakage while approximately preserving the long-tailed family-size distribution. Family-held-out performance was used for model selection because this setting tests transfer to previously unseen family patterns.

For precursor classification, negative examples were sampled from RNAcentral across non-miRNA RNA biotypes and supplemented with curated pseudo-hairpins. RNA secondary structures were predicted with RNAfold and encoded as positionally aligned dot-bracket strings. Sequence and structure channels were cropped synchronously during structure-constrained augmentation so that native stem pairing was preserved.

### miRstring architecture

miRstring formulates precursor-to-mature miRNA prediction as localization of the four boundaries defining the 5p and 3p mature segments. Nucleotide sequences were encoded by the pretrained ERNIE-RNA backbone [32], whereas aligned secondary structures were represented by a trainable structural embedding. A residual structure-guided cross-attention module used sequence representations as queries and structural representations as keys and values. For attention head h, the cross-modal update was computed as:

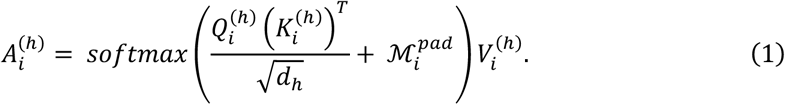

The structure-conditioned sequence states were then fused with the original structural embeddings and projected into a shared latent space:

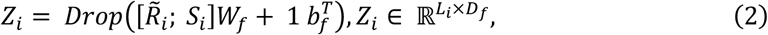

The fused states were processed by a shared Transformer encoder [28] to capture maturation cues common across taxa. The resulting representations were subsequently routed, using known superclass labels, to dedicated tail encoders for Metazoa, Viridiplantae, Viruses and Others, providing lineage-specific refinement without training independent models for each group.

### Boundary prediction and training objective

Clade-specific heads produced position-wise logits for the 5p start, 5p end, 3p start and 3p end coordinates. Padded positions were excluded from normalization. For boundary type k, the masked position probability was defined over the valid precursor coordinates Ωᵢ as:

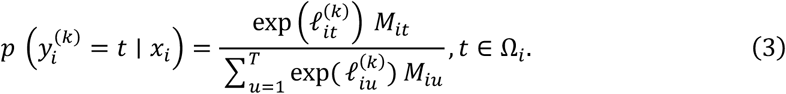

Boundary localization was optimized by the negative log-likelihood of all annotated 5p and 3p boundaries. With δᵢ⁵ and δᵢ³ indicating whether the corresponding mature arm was annotated, the boundary loss was:

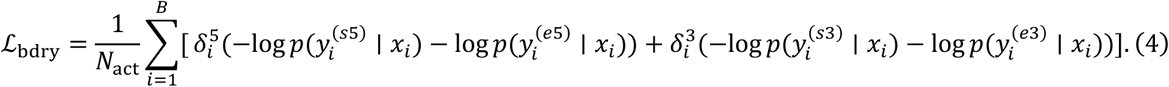

Two lightweight auxiliary classifiers were trained from frozen miRstring representations: a precursor-validity head distinguishing bona fide miRNA precursors from other RNAs and a guide-arm head assigning the recovered guide product to the 5p or 3p arm.

### Optimization and staged training

Models were optimized with AdamW [29] using a batch size of 32, automatic mixed precision, gradient clipping at 1.0, cosine warmup and early stopping. Training proceeded in four stages. First, the RNA language backbone was frozen while the remaining modules were trained on miRBase. The backbone was then unfrozen and jointly refined with the shared encoder using a smaller learning rate and balanced animal-plant mini-batches. Finally, the shared backbone was frozen and the Metazoa and Viridiplantae branches were refined separately, with span-level validation accuracy used for branch-specific model selection.

### Model-guided artificial miRNA scaffold selection

For each 21-nt artificial miRNA (amiRNA) guide, candidate constructs were generated from conserved endogenous precursor scaffolds. The designed guide replaced the endogenous guide sequence, paired passenger-arm positions were updated by Watson-Crick complementarity, and unpaired positions were initially retained. Passenger-arm bulges were then refined by enumerating substitutions. Each candidate was refolded with RNAfold, and variants were retained only when the predicted secondary structure matched the native scaffold fold. Candidates were prioritized by minimal folding-energy perturbation and the number of edited bulge positions.

The retained precursors were scored with miRstring at the expected guide boundaries. If sₐ and eₐ denote the expected start and end labels for the inserted guide on arm a, the guide-boundary confidence was:

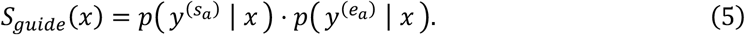

Constructs with unresolved guide-arm assignment or invalid boundary configurations were removed. High-confidence candidates were first selected within scaffold families and then ranked across families to preserve scaffold diversity for experimental screening.

### Experimental validation and statistics

Candidate 21-nt target sites against LUC and the endogenous Nicotiana benthamiana genes PDS and CHLH were designed using WMD3/Web MicroRNA Designer according to established plant amiRNA design principles [30,31]. Conventional MIR319-based constructs served as scaffold-reuse controls, whereas model-guided constructs used the two highest-scoring scaffold families for each target. Final precursor sequences were synthesized as DNA fragments and inserted into a pCAMBIA1303-derived binary vector.

For LUC validation, the reporter and amiRNA expression vector were co-infiltrated into one side of N. benthamiana leaves, while the opposite side received the LUC reporter alone as a paired internal control. For endogenous-gene validation, amiRNA constructs targeting PDS or CHLH were introduced by Agrobacterium tumefaciens-mediated transient expression, with empty-vector infiltrations as negative controls. Total RNA was isolated, reverse-transcribed and quantified by RT-qPCR using the primers listed in Supplementary Table 2. Target-gene expression was normalized to ACTIN. Relative expression was calculated using the ΔΔCt method:

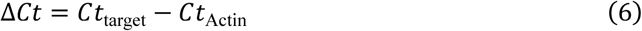

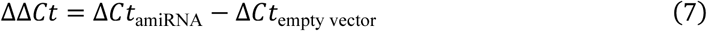

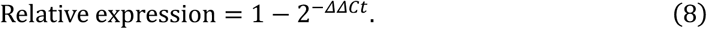

Statistical comparisons between model-guided constructs and the conventional MIR319 scaffold were performed on ΔCt values using one-sided unpaired Student’s t-tests.

## 3. Results

### Design overview of the miRstring architecture

miRstring takes as input a tokenized precursor sequence together with its RNA secondary structure and a species label that serves as an identifier for downstream clade-specific modules, and predicts the mature miRNA and miRNA* substring along the precursor hairpin **(Figure. 1)**. Spetically, miRstring predicts the four boundary sites of miRNA/miRNA* duplex, which are designated as the 5p start and 5p end on the 5p arm of the stem loop structure, and the 3p start and 3p end on the 3p arm of the stem loop.

**Figure. 1.**
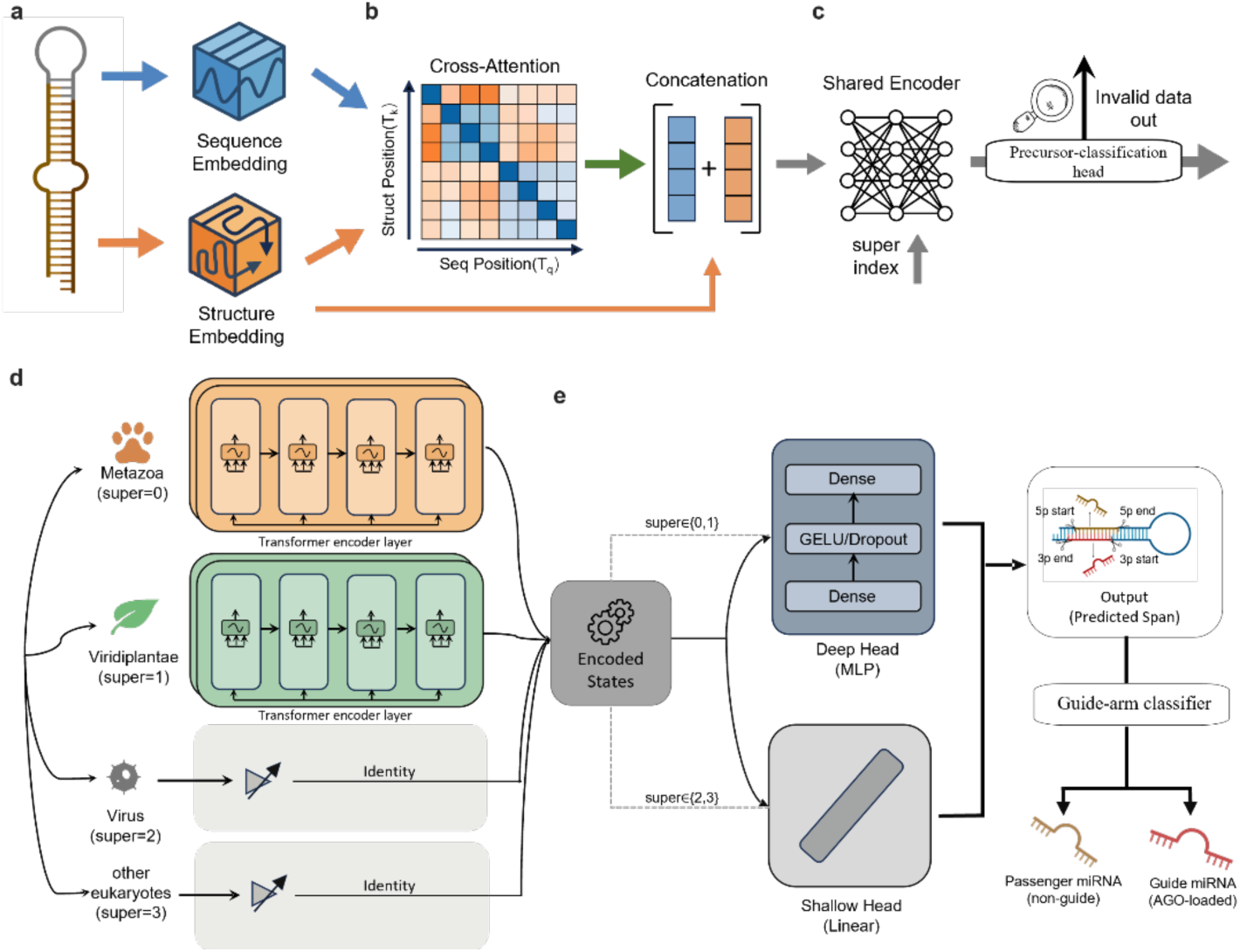
Overview of the miRstring architecture. **a,** The precursor hairpin is represented by two input modalities separately: sequence and RNA secondary structure. **b,** A structure-guided cross-attention module integrates sequence and structural information. **c,** The fused representation is processed by a shared encoder, together with a precursor-validity judging branch. **d,** Encoded states are routed through super-clade-specific modules to capture lineage-dependent processing patterns. **e,** Clade-adapted heads decode the shared representations and jointly localize the miRNA/miRNA* duplex within the precursor by predicting its four defining boundaries, which are designated as 5p start and 5p end on the 5p arm of precursor hairpin (stem loop), and 3p start and 3p end on the 3p arm of given stem loop.

Sequence and structure are first encoded separately using a pretrained RNA language model (ERNIE-RNA) and a trainable structural embedding layer, respectively **(Figure. 1a)**[32]. Structure is then integrated at two complementary stages: first through a structure-guided cross-attention block, in which sequence representations act as queries and projected structural encodings serve as keys and values, and then through explicit fusion with the original structure embeddings, so that structural signals remain available throughout downstream processing **(Figure. 1b)**.

After sequence–structure integration, miRstring uses a shared Transformer encoder to learn shared boundary-associated cues that recur across diverse precursors **(Figure. 1c)**. In parallel, we attach a lightweight sequence-level Binary precursor-classification head to the shared representation to estimate whether an input is likely to correspond to a bona fide miRNA-producing precursor, providing a global precursor-validity judge. Although the detailed biogenesis and targeting logic differs between major groups, miRNA production broadly follows a common scaffold: hairpin transcripts are processed by RNase III–type enzymes into a duplex and loaded into Argonaute effector complexes, which motivates learning a unified representation that captures boundary-adjacent sequence–structure regularities shared across taxa[33,34].

To account for systematic differences across broad groups, we introduce super-clade-routed tail encoders for Metazoa, Viridiplantae, Viruses and Others **(Figure. 1d)**. These tails provide post-encoder specialization that refines the shared representation with clade-specific processing context without fragmenting the entire model into separate networks. In bilaterian animals, target recognition is dominated by the 5′ seed region, such that even a 1-nt shift at the miRNA 5′ end can substantially alter targeting specificity[3]. In plants, by contrast, target recognition typically depends on extensive miRNA–target complementarity across much of the guide and often leads to AGO-mediated slicing opposite guide positions 10–11[4,5]. These distinct targeting regimes suggest that the functional consequences of duplex boundary placement are expressed differently across clades, motivating clade-aware refinement after a shared encoder [3,4]. Routing to tails is implemented as hard assignment using the known super-clade label. This keeps specialization explicit and stable, avoiding the added optimization complexity of jointly learning mixture weights with boundary localization, and ensuring that any downstream differences arise from the designated clade-specific tail.

Finally, a clade-specific head bank maps tail representations to boundary logits, with shallow multilayer perceptron (MLP) heads used for the better-annotated Metazoa and Viridiplantae groups and linear heads used for the smaller or noisier Viruses and Others groups to limit overfitting **(Figure. 1e)**. miRstring then predicts the mature-miRNA duplex through masked position classification over the precursor, yielding four boundary-specific score tracks for precise cut-site localization. We further introduced an auxiliary guide-arm classifier to determine, for each recovered duplex, which arm functions as the guide strand and which as the passenger strand, using database annotations as supervision.

### Multi-stage training of miRstring on curated miRNA precursor resources

To construct a unified precursor-centred resource, we integrated miRNA annotations from miRBase, MirGeneDB and PmiREN[35–37]. Restricting the collection to high quality precursors with annotated mature products yielded a final dataset of 77,708 precursors spanning 414 species **(Figure. 2a)**. This integration substantially expanded species coverage beyond miRBase, with MirGeneDB contributing 55 additional metazoan species and PmiREN contributing 88 further species **(Figure. 2b)**; among these newly added species, we reserved 25 from the more stringently curated MirGeneDB and 10 from PmiREN as strict species-held-out evaluation sets to assess performance in the scenario of miRNA identification from previously unseen species.

**Figure. 2.**
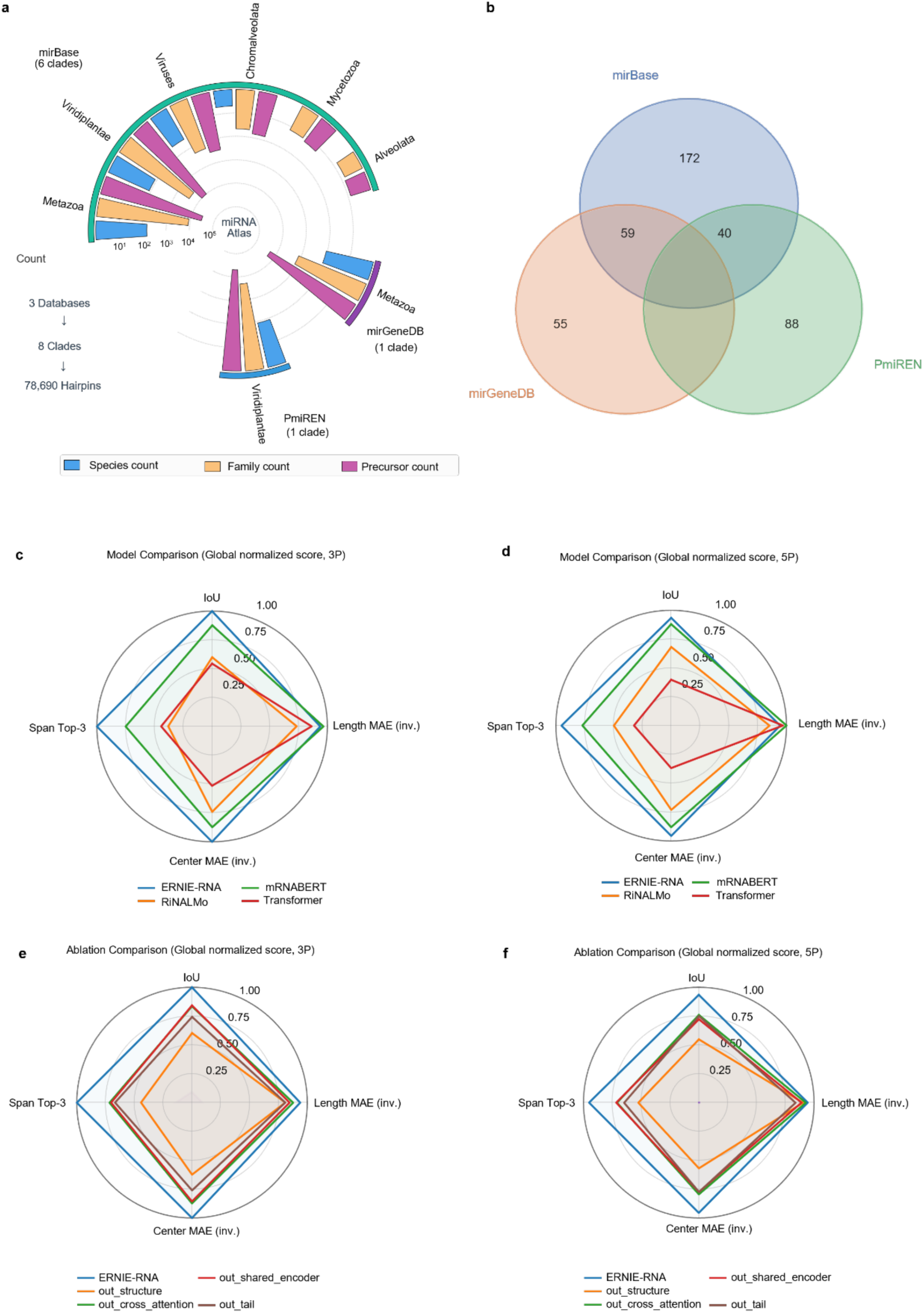
Dataset composition of miRNA precursors and ablation study of miRstring. **a,** Overview of the curated miRNA precursor dataset assembled from miRBase, MirGeneDB and PmiREN across major taxonomic clades. Bars indicate species count, miRNA-family count and precursor count. **b,** Species-level overlap among miRBase, MirGeneDB and PmiREN, showing database-specific and shared species coverage. **c,d,** Benchmark of miRNA/miRNA* sequence-representation from the 3p arm and 5p arm of precursors, respectively. The downstream miRstring architecture was kept fixed while the sequence-representation module was replaced by ERNIE-RNA, mRNABERT, RiNALMo or a Transformer-based encoder. **e,f,** Component ablation analysis of miRNA/miRNA* sequence-representation from the 3p arm and 5p arm of precursors, respectively. Ablated variants removed the structural input channel, the structure-guided cross-attention module, the shared contextual encoder or the super-clade-specific tail encoder from the full ERNIE-RNA-based model. Radar axes summarize span-level localization performance using IoU, Span Top-3 accuracy, centre MAE and length MAE. Scores were globally normalized to 0–1 for visualization; IoU and Span Top-3 were normalized directly, whereas centre and length MAE were inversely normalized so that higher values indicate smaller errors and better overall performance.

For model selection, we used a strict family-held-out protocol, with all precursors from the same miRNA family assigned to the same split. Because miRNA families are organized around closely related mature sequences and often recur across species, species-only splits can remain permissive, whereas family isolation provides a stricter test of transfer to unseen family patterns[3,36]. Family-held-out splits were generated in a size-stratified manner to account for the long-tailed family-size distribution; full details are provided in Methods.

To train the miRNA precursor-classification head, we constructed a complementary binary dataset whose negative set was sampled from RNAcentral across diverse non-miRNA RNA biotypes, including transfer RNA (tRNA), ribosomal RNA (rRNA), small nuclear RNA (snRNA), small nucleolar RNA (snoRNA), other non-coding RNA (ncRNA) and messenger RNA (mRNA), and further supplemented with curated pseudo-hairpin negatives **(Supplementary Figure. 1a)**[38].

Ablation analysis further supported the overall design of miRstring **(Figure. 2c-2f)**. For predicting miRNA/miRNA* sequence representation from 5p arm and 3p arm of precursor stem loop, the full model consistently achieved the strongest overall performance, indicating that the complete architecture was optimal for duplex-boundary recovery. Among the tested sequence encoders, ERNIE-RNA provided the best results, outperforming the non-pretrained and alternative pretrained encoders (RiNALMo and mRNABERT)[32,39,40], and thus yielding the strongest sequence representation for downstream localization **(Figure. 2c,d)**. Beyond encoder choice, removal of any individual input or architectural component reduced performance to varying degrees, showing that structure-aware inputs, cross-modal integration and routed downstream decoding each made non-redundant contributions **(Figure. 2e,f)**. Taken together, these ablations support a cooperative design in which high-quality sequence representations, explicit structural information and clade-aware decoding act together to enable accurate mature-miRNA span recovery.

### Evaluation of span-level boundary recovery and cross-species generalization

We first evaluated miRstring at nucleotide resolution on the miRNA precursor family-held-out validation sets using signed errors for the four mature-miRNA boundaries, defined as predicted minus annotated position. To prevent large families from dominating the analysis, errors were averaged within each family before visualization. Across both Metazoa and Viridiplantae, all four boundary types remained centred close to zero, indicating little systematic directional bias in boundary placement **(Figure. 3a,b)**. The remaining spread revealed heterogeneous difficulty across held-out family contexts, with broader dispersion in Viridiplantae than in Metazoa.

**Figure. 3.**
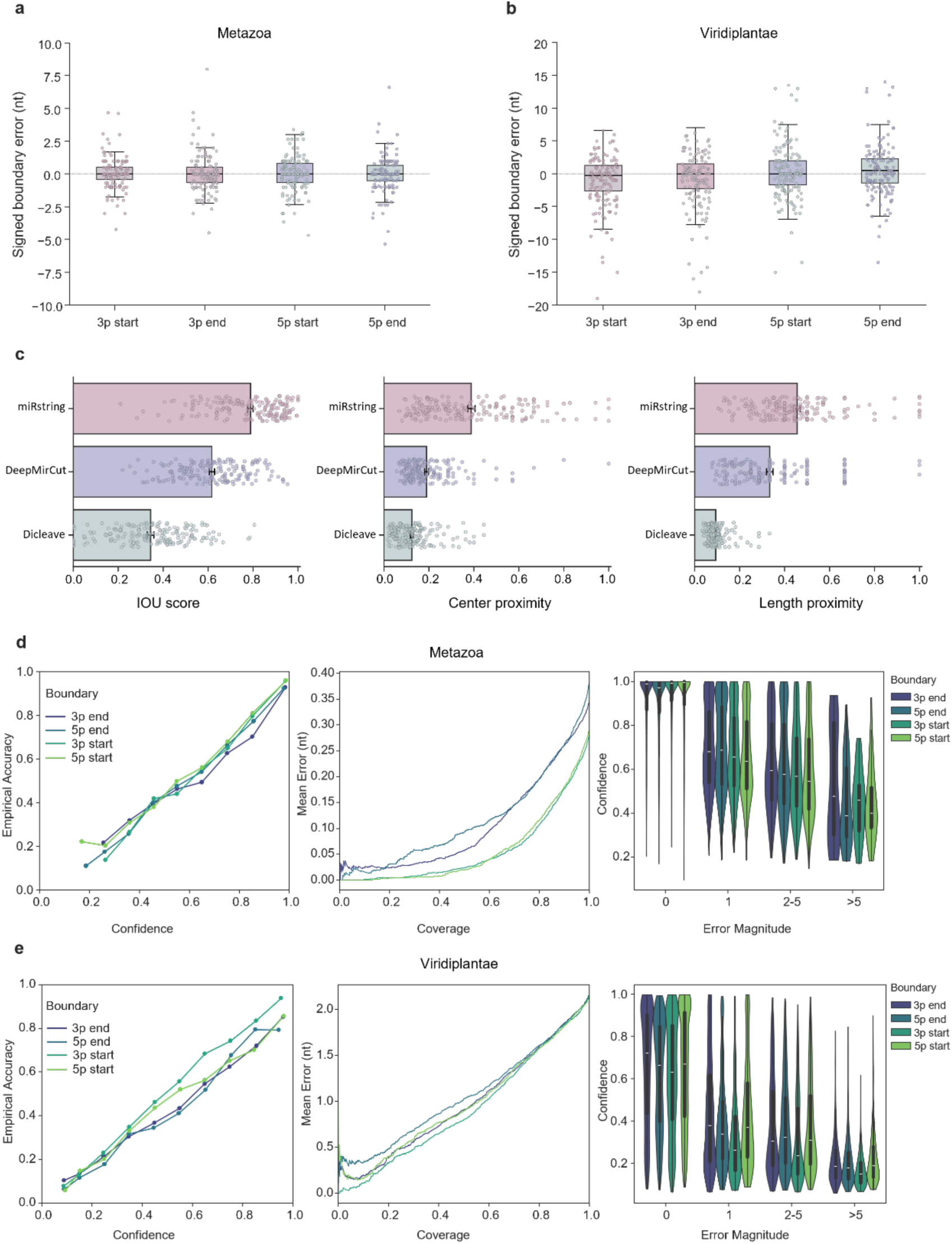
miRNA Boundary error distributions, miRstring benchmark comparison and confidence calibration across miRNA family-level and species-level holdout sets. **a,** Distribution of miRNA family-averaged signed boundary errors for the four miRNA/miRNA* boundary types (5p start, 5p arm end, 3p arm start and 3p arm end) in metazoa, evaluated on the validation set comprising families not seen during training. Signed boundary error was defined as predicted boundary position minus true boundary position (nt). Each point represents one family, with errors averaged within family before plotting. Boxes indicate the interquartile range (IQR), centre lines indicate the median, and whiskers extend to 1.5 × IQR. **b,** Same analysis as in a, shown for viridiplantae on the miRNA family-level holdout validation set. **c,** miRNA family-level performance comparison of miRstring, DeepMirCut and DiCleave on the held-out validation set comprising families not seen during training. For each miRNA family, performance was summarized using intersection over union (IOU), centre proximity and length proximity, where proximity was defined as 1 / (1 + MAE), such that higher values indicate better performance. Bars denote the mean across families and points denote individual families. Results of DeepMirCut and DiCleave are reported after training on the corresponding non-overlapping training data and evaluation on the held-out metazoa species set. **d,** Uncertainty diagnostics for the four miRNA/miRNA* boundary heads (5p start, 5p end, 3p start and 3p end) on the held-out metazoa species set, including reliability diagrams, risk–coverage curves and confidence distributions stratified by absolute boundary error. Lower boundary error is associated with higher predicted confidence, and confidence-based selection reduces mean boundary error at matched coverage. **e,** Same analysis as in d, shown for the held-out viridiplantae species set.

We next compared miRstring with DeepMirCut and DiCleave on the same family-held-out benchmark[22,23]. Performance was summarized at the miRNA-family level using Intersection over Union (IoU), center mean absolute error (MAE) and length MAE, with center and length errors additionally transformed to bounded proximity scores, 1/(1 + MAE), for visualization **(Figure. 3c)**. Across all metrics, miRstring consistently outperformed both baselines, achieving a mean family-level IoU of 0.789, a center MAE of 2.664 nt and a length MAE of 1.590 nt. Relative to DeepMirCut and DiCleave, miRstring reduced center MAE by 53.0% and 77.3%, respectively, while also improving IoU and reducing length MAE, indicating more faithful recovery of both the location and extent of mature-miRNA duplexes.

To assess generalization under species shift, we further evaluated miRstring on strictly held-out species using reliability diagrams, risk–coverage curves and confidence distributions stratified by error magnitude **(Figure. 3d,e)**. Across both clades, confidence tracked prediction quality well: higher-confidence predictions showed higher empirical accuracy, lower mean boundary error and a clear enrichment of near-exact boundary calls, whereas larger errors were concentrated at lower confidence. Although absolute error levels differed between Metazoa and Viridiplantae, this confidence–quality relationship was preserved in both groups, supporting the use of miRstring uncertainty estimates for selective prediction and quality control in unseen species.

Beyond uncertainty calibration, we examined signed boundary errors on the species-held-out sets, averaged within each species **(Figure. 4a,b)**. Across both Metazoa and Viridiplantae, all four duplex boundaries remained centred near zero, indicating minimal systematic bias under true species shift, although some between-species variation remained.

**Figure. 4.**
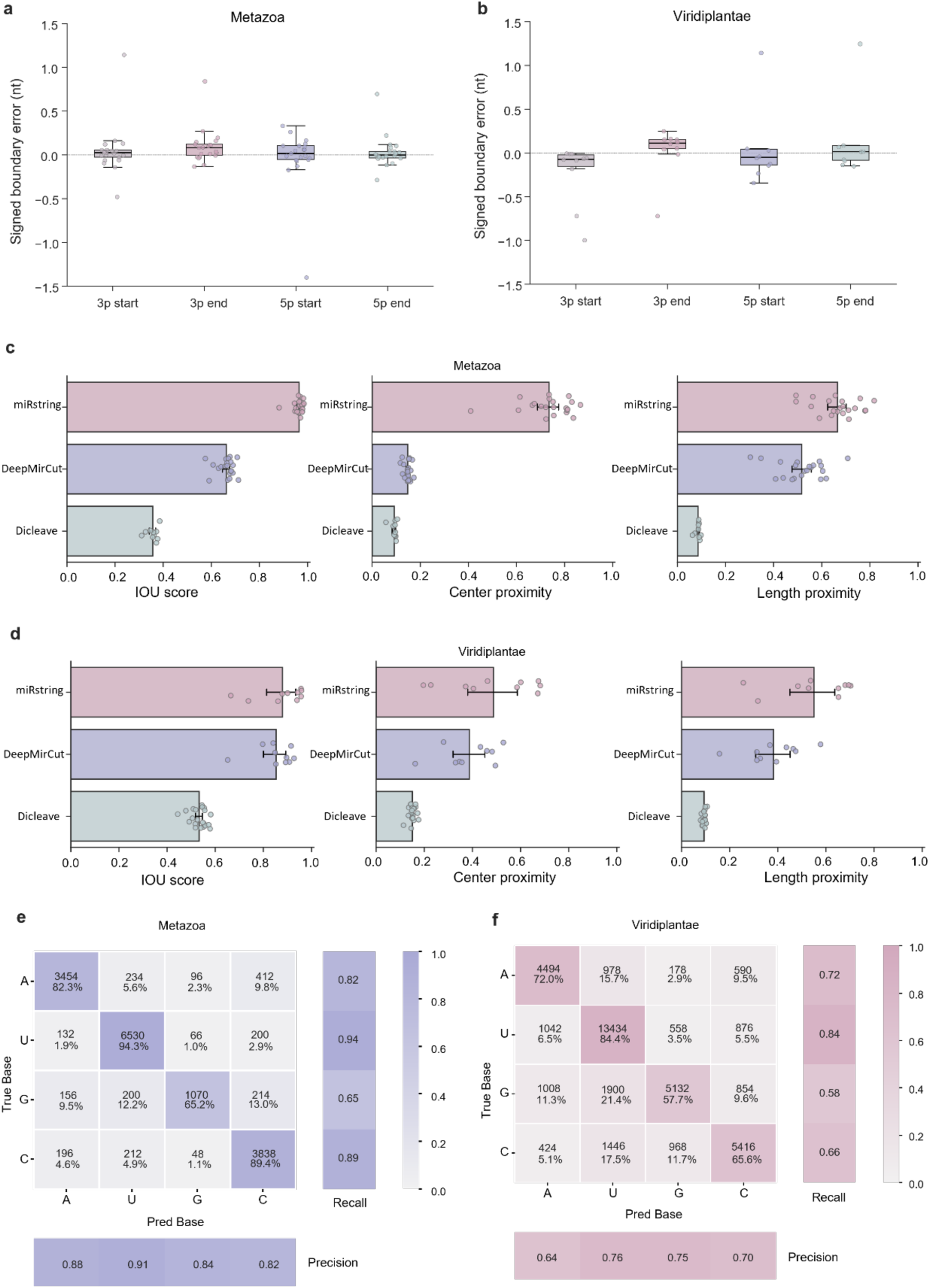
Species-averaged signed miRNA/miRNA* boundary errors on held-out family-level validation sets. **a,** Distribution of species-averaged signed boundary errors for the four miRNA/miRNA* boundary types (3p start, 3p end, 5p start and 5p end) in metazoa, evaluated on the held-out validation set comprising families not seen during training. Signed boundary error was defined as predicted boundary position minus true boundary position (nt). Each point represents one species, with boundary errors averaged within species before plotting. Boxes indicate the interquartile range (IQR), centre lines indicate the median, and whiskers extend to 1.5 × IQR. **b,** Same analysis as in a, shown for viridiplantae on the held-out family-level validation set. Positive values indicate overestimation of boundary position, whereas negative values indicate underestimation. For visualization, points outside the displayed y-axis range were omitted, whereas boxplot statistics were computed from all species-level values. **c,** Species-level performance comparison of miRstring, DeepMirCut and DiCleave on the held-out metazoa species set, comprising species excluded from training. Results of DeepMirCut and DiCleave are reported after training on the corresponding non-overlapping training data and evaluation on the held-out metazoa species set. **d,** Same analysis as in c, shown for the held-out viridiplantae species set. **e,** Confusion matrix of first-base prediction of miRNAs on the held-out metazoa species set. Rows represent true bases and columns represent predicted bases. The main matrix is row-normalized, with each cell showing the number of observations and the corresponding percentage within the true-base class. The right-hand and bottom panels report recall and precision, respectively. **f,** Same analysis as in e, shown for the held-out viridiplantae species set.

We then compared miRstring with DeepMirCut and DiCleave on the strictly species-held-out sets at the species level **(Figure. 4c,d)**[22,23]. In both held-out metazoan and plant species, miRstring ranked best across IoU, center MAE and length MAE, with especially marked gains in metazoans. These results show that the advantages of miRstring extend beyond held-out families to genuinely unseen species.

Notably, performance on the species-held-out benchmarks exceeded that on the family-held-out benchmark, with mean IoU increasing from 0.789 at the family level to 0.965 in held-out metazoan species and 0.880 in held-out plant species, while center MAE decreased from 2.664 nt to 0.396 nt and 1.464 nt, respectively. This gap is consistent with family-held-out evaluation being the stricter test of generalization: species isolation can still preserve familiar family-level signatures, whereas family isolation requires transfer to genuinely unseen family patterns.

To assess whether miRstring recapitulates biologically meaningful first-nucleotide preferences in unseen species, we evaluated first-base predictions using confusion matrices. In plants, where most miRNAs carry a 5′ uridine and are preferentially sorted into AGO1.

miRstring recovered U particularly well while still distinguishing A, G and C, rather than collapsing to the dominant class **(Figure. 4f)**[41]. A similar pattern was observed in Metazoa: despite weaker determinism, metazoan miRNAs are also enriched for 5′ U, and miRstring preserved this expected distribution on held-out species without losing discrimination among non-U nucleotides **(Figure. 4e)**[42]. These results indicate that the model generalizes biologically meaningful 5′-nucleotide preferences under species shift.

Additionally, the lightweight binary classifier, trained on the shared representation, achieved strong discrimination of miRNA precursors from other RNAs, with a validation area under the receiver operating characteristic curve (AUROC) of 0.9975 and a precision-recall area under the curve (PR-AUC) of 0.9984 **(Supplementary Figure. 1b-d)**. Furthermore, the auxiliary classifier for recognizing guide miRNAs from miRNA/miRNA* duplex also performed well, reaching a validation AUROC of 0.8988 and a PR-AUC of 0.9186 **(Supplementary Figure. 2)**.

### Attention-guided representations of miRNA boundary microenvironments

We next leveraged self-attention maps from the shared encoder to examine which precursor positions were most informative for duplex-boundary localization. In representative precursors, attention was enriched mainly along the stem and clustered near the four boundaries defining the miRNA duplex, rather than across unpaired loops or flanking regions. miRstring has learned to prioritize boundary-adjacent sequence–structure context as the key local determinant of Dicer cleavage-site definition **(Figure. 5a, b)**. To test whether these patterns generalized beyond representative examples, we quantified attention enrichment in a random subset of validation precursors and compared the results with null expectations. Attention was depleted at unpaired positions, enriched at paired stem positions, and strongest in boundary-proximal regions **(Figure. 5c)**, with mean attention importance decreasing as distance from the nearest duplex boundary increased **(Figure. 5d)**.

**Figure. 5.**
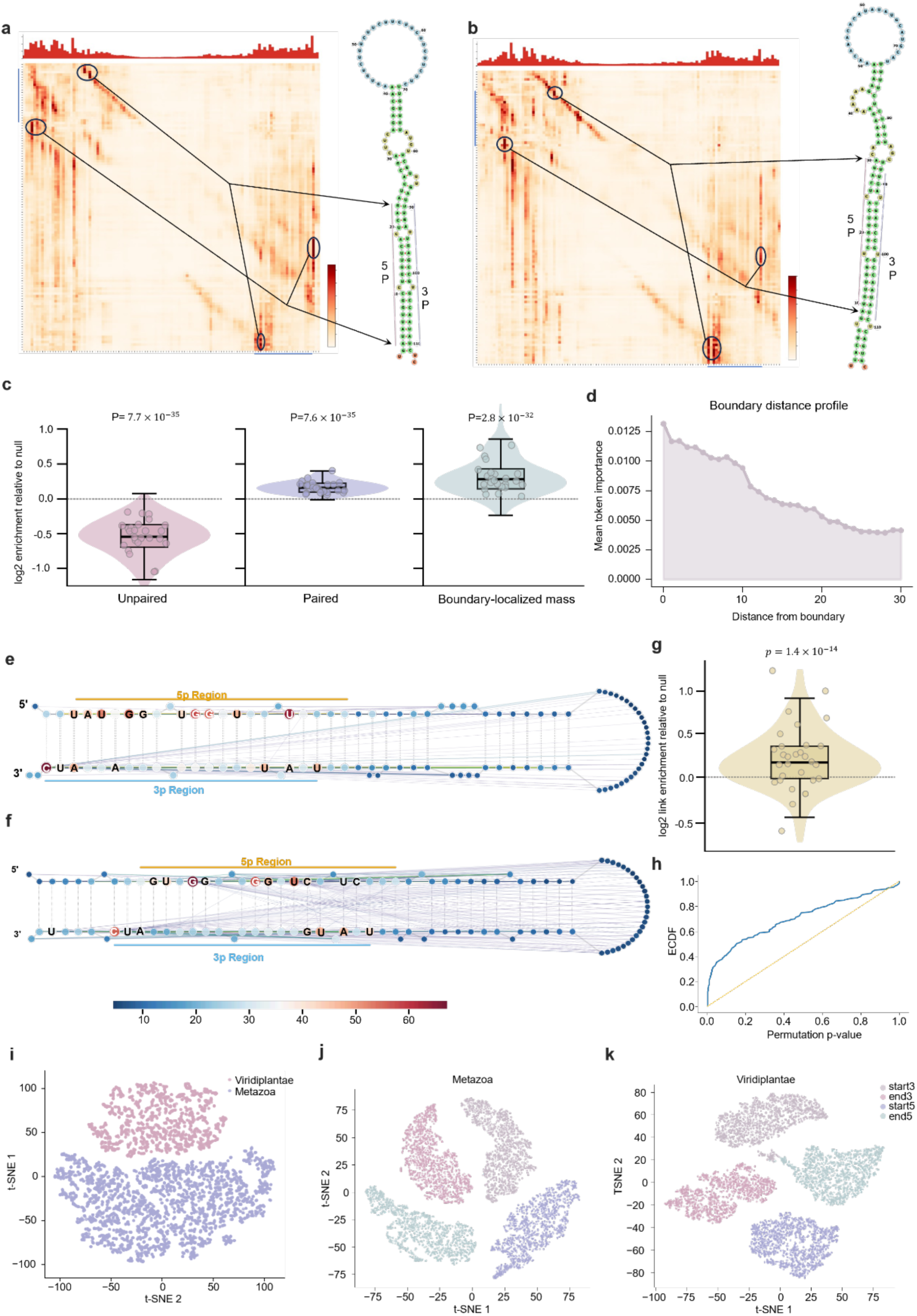
Learned attention highlights miRNA boundary-proximal sequence–structure cues. **a,b,** Representative self-attention maps from the shared encoder, shown as query × key positions along the precursor sequence. Warmer colours indicate stronger attention weights, illustrating how the model aggregates contextual evidence across the miRNA precursor. Highlighted attention hotspots are linked by arrows to their corresponding positions on the predicted hairpin secondary structure, showing that the learned interactions align with structurally related regions of the stem– loop. **c,** Distributions of log2 attention enrichment relative to a null expectation for unpaired positions, paired positions and boundary-localized mass within ±2 nt of the mature-miRNA boundaries across 200 examples. Dashed lines indicate zero enrichment. Violins show the full distribution, and boxes indicate the median and interquartile range (IQR), both computed from all 200 examples. For visualization only, 30 randomly sampled examples are shown as jittered points. P values were calculated using one-sided Wilcoxon signed-rank tests against 0. **d,** Boundary distance profile of mean attention importance. Mean attention importance is plotted as a function of distance to the nearest miRNA/miRNA* boundaries. **e,f,** Structure-mapped attention landscapes for the same two representative precursors shown in d and e, respectively. Nucleotides are arranged according to the predicted hairpin secondary structure. Node colour and size indicate attention-derived importance scores, whereas overlaid curves represent high-weight attention links whose values exceed the 98th percentile of the full attention matrix for each example. Coloured bars mark the predicted mature-miRNA regions. **g,** Distribution of log2 enrichment relative to a null expectation for boundary-centred wiring across 200 examples. The dashed line indicates zero enrichment, corresponding to no excess boundary-centred wiring relative to the null. Violins show the full distribution, and overlaid boxplots indicate the median and interquartile range (IQR), both computed from all 200 examples. For visualization only, 30 examples were randomly sampled and displayed as jittered points. Statistical significance was assessed using a one-sided Wilcoxon signed-rank test against 0, testing whether boundary-centred wiring enrichment was consistently positive across examples. **h,** Empirical cumulative distribution function (ECDF) of permutation-derived right-tailed P values for boundary-centred wiring enrichment across the same 200 examples used in g. For each example, the P value quantifies the probability, under a permutation-based null model estimated from 500 permutations, of observing an enrichment statistic at least as large as the one measured in the real attention graph. The dashed diagonal indicates the uniform expectation under the null. Upward deviation of the ECDF above the diagonal indicates an excess of small P values, supporting consistent boundary-centred wiring enrichment across examples. **i,** t-SNE projection of learned latent representations for Viridiplantae and Metazoa. Each point represents one sample in the latent space and is coloured by clade, showing separation between viridiplantae and metazoa precursor representations in the learned feature space. **j-k,** Two-dimensional projection of latent representations from Metazoa (j) and Viridiplantae(k), coloured by boundary type (3p start, 3p end, 5p start and 5p end). Each point represents one sample in the learned feature space. The embedding shows separation among boundary-specific representations within precursors.

On examining these attention-enriched boundary regions more closely, we found that they frequently exhibited local sequence–structure transitions, including paired-to-unpaired shifts and small bulges **(Supplementary Figure. 3a)**, as well as distinct motif enrichments around the corresponding cleavage sites **(Supplementary Figure. 3c)**. This is biologically plausible, as prior mechanistic studies have shown that local bulges, mismatches and other boundary-proximal structural features can influence Microprocessor recognition and Dicer cleavage accuracy in certain cases[30,43–45], suggesting that the boundary microenvironments captured by miRstring may encode informative signals for enzymatic cleavage-site recognition.

To move beyond marginal attention hotspots and examine how the model encodes relationships between different positions along the precursor, we next projected shared-encoder attention onto the hairpin secondary-structure layout and visualized sparse high-weight links between nucleotide positions **(Figure. 5e,f)**. In this view, the strongest interactions were not diffusely distributed across the precursor, but instead concentrated around a limited number of local hubs near the mature-duplex boundaries, indicating preferential coupling around cleavage-relevant regions. Permutation-based analysis confirmed that this boundary-centred wiring was consistently enriched across the validation subset **(Figure. 5g, h)**.

Notably, we observed that the strongest pairwise coupling occurred between the start and end boundaries of the same mature miRNA, both for the 3’ segment and for the 5’ segment **(Supplementary Figure. 3b)**. Because a mature miRNA is jointly defined by its start and end cleavage boundaries, this pattern suggests that miRstring jointly encodes the two boundaries of the same mature product, rather than representing them independently, thereby learning mature-miRNA segment integrity.

Two-dimensional projection of the learned feature space separated Metazoa from Viridiplantae at the precursor level **(Figure. 5i)**, indicating that the learned representation captures clade-dependent sequence–structure characteristics, within each clade further resolved the four duplex-boundary types, indicating that distinct cleavage positions are encoded with distinct local contextual features **(Figure. 5j, k)**. To relate the model’s learning signal to evolutionary novelty, we mapped each miRNA family to an inferred origin on a MirGeneDB species phylogeny and quantified its training influence under a strict family-heldout protocol. We observed lineage-structured influence patterns: families originating in different clades contribute unevenly to heldout-family generalization, with certain terminal lineages showing concentrated high-impact families. This suggests that beyond broadly shared biogenesis cues, lineage-specific family innovations provide distinct gradient signals that shape how the model resolves boundary localization in unseen families **(Supplementary Figure. 4)**. Taken together, these results suggest that the model spontaneously reconstructs the multiscale constraints imposed on the stem–loop during miRNA processing.

### The amiRNA design enables sequence-specific scaffold selection

Previous studies have employed a limited number of miRNA precursors as scaffolds for artificial miRNA (amiRNA). For instance, miR319 percussor is widely used for scaffold of amiRNA in Arabidopsis[30]. Using miRstring, we aimed to develop a scaffold-aware, model-guided pipeline for artificial miRNA construction that preserves native precursor context while supporting sequence-specific scaffold selection **(Figure. 6)**. Starting from curated conserved scaffolds from Metazoa or Viridiplantae, we graft user-specified guide sequences into the duplex guide window and update only the paired counterparts required to maintain stem consistency **(Figure. 6a)**. We then refine the passenger side by enumerating substitutions at bulge positions, retaining only structure-preserving variants and selecting the best candidate for each scaffold–guide pair based on minimal energetic and structural perturbation. Finally, generated constructs are scored with miRstring as a boundary-signature oracle, and high-confidence candidates are prioritized while preserving scaffold diversity **(Figure. 6b-c)**. Together, this pipeline enables scalable scaffold-aware artificial miRNA design and prioritization for downstream experimental screening.

**Figure. 6.**
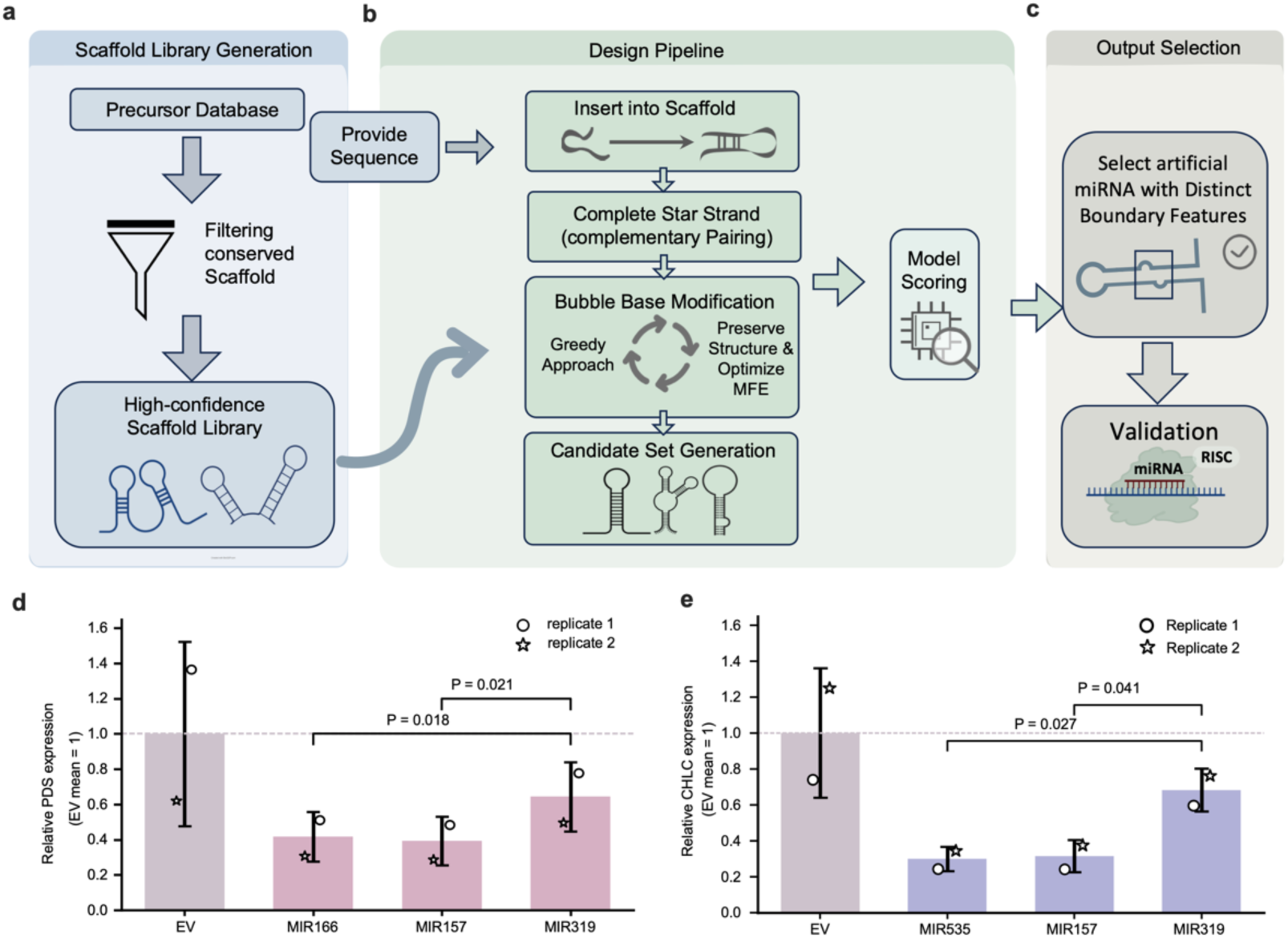
Scaffold-aware, model-guided pipeline for artificial miRNA design. **a,** Construction of a high-confidence scaffold library from conserved precursor backbones. **b,** Sequence-specific design workflow: guide insertion, duplex completion, local bulge refinement and miRstring-based scoring of candidate constructs. **c,** Selection of high-confidence artificial miRNA candidates with distinct boundary features for downstream validation. **d,** Relative expression of endogenous *PDS* in *Nicotiana benthamiana* leaves transiently expressing the indicated amiRNA constructs or the empty-vector control. **e,** Relative expression of endogenous *CHLH* under the same transient-expression assay. Target-gene expression was normalized to *ACTIN* and expressed relative to the mean empty-vector control, which was set to 1. Bars show mean ± s.d. from two independent biological replicates; individual points indicate biological replicates, with different marker shapes denoting replicate 1 and replicate 2. Statistical comparisons were performed on ΔCt values using one-sided unpaired Student’s *t*-tests comparing model-guided constructs with the conventional MIR319 scaffold; displayed *P* values indicate the tested comparisons.

To test whether the model-guided amiRNA designs could mediate functional repression in planta, we established a paired transient luciferase (LUC) reporter assay in *Nicotiana benthamiana (N. benthamiana)* leaves. For each construct, the LUC reporter and amiRNA expression vector were co-infiltrated into one side of the leaf, whereas the opposite side received the LUC reporter alone as an internal paired control. We further tested two independent target sites within the LUC reporter to assess repression across distinct target contexts. Representative bioluminescence images showed that, across multiple amiRNA constructs, including the two top-scoring model-guided scaffolds for each target site and the canonical MIR319 scaffold, the LUC signal was repressed by amiRNA compared to the control **(Supplementary Figure. 5a,b)**, This result indicates that these scaffolds can produce amiRNA to functionally repress its target reporter in plant cells.

To obtain a quantitative transcript-level readout, we further tested the selected amiRNA constructs against two endogenous pigmentation-related target genes, *PDS* and *CHLH*, using transient expression followed by reverse transcription quantitative polymerase chain reaction (RT-qPCR) in *N. benthamiana* leaves. Target-gene expression was normalized to Actin and compared with the empty-vector control. Overall, the selected scaffolds produced substantial reductions in endogenous target-gene expression **(Figure. 6e,f)**. In the amiRNA-targeted *PDS* assay, the model-selected MIR166 and MIR157 scaffolds reduced *PDS* transcript abundance by ∼60%, displaying significantly higher efficiency than the conventional MIR319 scaffold (∼40%). Similarly, in the *CHLH* assay, the model-selected MIR535 and MIR157 scaffold significantly decreased *CHLH* transcript abundance by ∼70%, indicating a stronger transcriptional repression effect than the conventional MIR319 scaffold (36.2%). Overall, the miRstring model can be used for scaffold selection to enhance the processing efficiency of rationally designed artificial miRNAs.

## 4. Discussion and conclusion

In this work, we present miRstring, a framework for recovering miRNA duplexes directly from precursor sequence and secondary structure under strict family-held-out and species-held-out evaluation. Our results show that this task remains tractable even under stringent out-of-distribution settings, and that miRNA duplex definition can be learned from precursor context rather than inferred only through curator- and experiment-intensive workflows.

miRstring was deliberately designed around the organizing principles of miRNA biogenesis. Through ablation, we found that its components worked best in combination, indicating that the model captures key information and shared regularities underlying precursor-to-duplex definition that support robust generalization across unseen families and species, and that provide useful clues for future studies of mature-miRNA prediction and processing. The same modeling strategy can be extended to additional biological groups with distinct small-RNA processing characteristics, including viral miRNAs as well as miRNA or miRNA-like pathways in protists, fungi and algae, as curated precursor annotations become available in these groups. Further decomposition of multi-head attention showed progressive enrichment from stem regions towards Dicer cleavage-proximal local neighborhoods. Given that local motif biases, paired-to-unpaired transitions and bulges have been implicated in the efficiency and precision of miRNA processing, these attention patterns may help prioritize candidate sequence–structure determinants for further studies of cleavage-site choice by miRNA processing machinery.

Notably, the performance gap between family-held-out and species-held-out evaluation led us to use the more stringent family-held-out setting for model selection, thereby favoring models that generalize beyond familiar family-level signatures. This is particularly relevant in miRNA annotation because family structure captures mature-sequence relatedness and, in curated resources, often shared evolutionary origin, while newly sampled lineages frequently introduce young, lineage-restricted or previously underrepresented families. The resulting model is therefore more relevant to mature-miRNA definition in newly sequenced or sparsely annotated settings, where previously unseen family contexts are common.

miRstring can be used to infer mature products directly from precursor context in newly sequenced or sparsely annotated genomes. Although experimental validation remains essential, especially for low-confidence or non-canonical cases, boundary localization with useful uncertainty estimates can still support annotation refinement and confidence-guided curation. miRstring also provides a new model-guided route for artificial miRNA design. Instead of relying only on a small set of generic empirical backbones, our scaffold-aware pipeline enables sequence-specific scaffold choice for diverse user-specified mature sequences while preserving precursor context as much as possible. In our experimental validation, model-guided constructs showed more stable expression and stronger knockdown than conventional scaffold-reuse designs. This provides a practical way to compare candidate scaffolds and prioritize designs before downstream experimental testing.

Importantly, several limitations of this work should be noted, which also point to directions for future research. One concerns structural representation: the current framework relies on a single predicted secondary structure per precursor, whereas real precursor RNAs may adopt alternative conformations that are not captured by a single minimum-free-energy fold. Incorporating structure ensembles or experimentally constrained folding information could therefore improve robustness in cases of conformational heterogeneity[46]. A second limitation lies in the supervision itself.

Although curated mature-miRNA annotations provide the scale needed for learning, they remain heterogeneous in confidence and incompletely capture context-dependent processing variation, including arm switching, shifted cleavage and microRNA isoform diversity[47,48]. Extending the training signal to richer small-RNA evidence and more explicit representations of cleavage variability may help bridge this gap. More broadly, current training resources remain uneven across clades, with viruses and other non-canonical groups still much more sparsely represented than metazoan and plant precursors. This limits how far the present framework can be generalized across the full diversity of miRNA biogenesis. Finally, although our artificial-miRNA design results support the practical utility of model-guided scaffold selection, broader validation across additional scaffolds, guide sequences and biological systems will be needed to determine how general this strategy is in practice. Together, these limitations define the clearest next steps for extending miRstring towards more structurally realistic, biologically comprehensive and experimentally grounded modeling. Taken together, our study suggests that miRNA duplex can be recovered from precursor context with substantial accuracy even under stringent out-of-distribution settings when sequence, structure and lineage-dependent processing context are modeled jointly. Beyond prediction itself, this framework supports more principled artificial-miRNA design. As curated small-RNA resources continue to expand, this work can support mature guide miRNA definition across newly sampled lineages and provide a practical basis for connecting precursor candidates to downstream functional investigation. More broadly, our study suggests an RNA language framework for model design and evaluation in biologically heterogeneous prediction tasks, in which meaningful generalization depends on aligning task definition, model architecture and evaluation strategy with domain-specific biological structure.

## 5. Acknowledgements

This work was funded by grants from National Natural Science Foundation of China (Grant No. 32370587) and Shanghai Municipal Education Commission (No. 2024AIYB005).

## 7. Data Availability Statement

The source code and pretrained model files for miRstring are archived in Zenodo at https://doi.org/10.5281/zenodo.21324293 and are also available from the corresponding GitHub release at https://github.com/ppksjjtu/miRstring/releases/tag/v1.0.0. The archived release includes the analysis and inference scripts, example input files, scaffold files, environment specification, pretrained checkpoint archive, and checksum file required to reproduce the software release described in this study.

## 8. Contributions

X.Y., and R.K.P. designed this study. R.K.P. and Y.F.T. performed the computational analysis. R.K.P., and X. L. performed the validation experiments. R.K.P., Y.F.T. and X.Y. wrote the manuscript. All authors reviewed and approved the final version of the manuscript.

