## Supplementary material for "miRstring: An RNA language model enables mature miRNA decoding and artificial small RNA design across species": Figure S (from another version of this preprint: version 1, 2026-09-20)

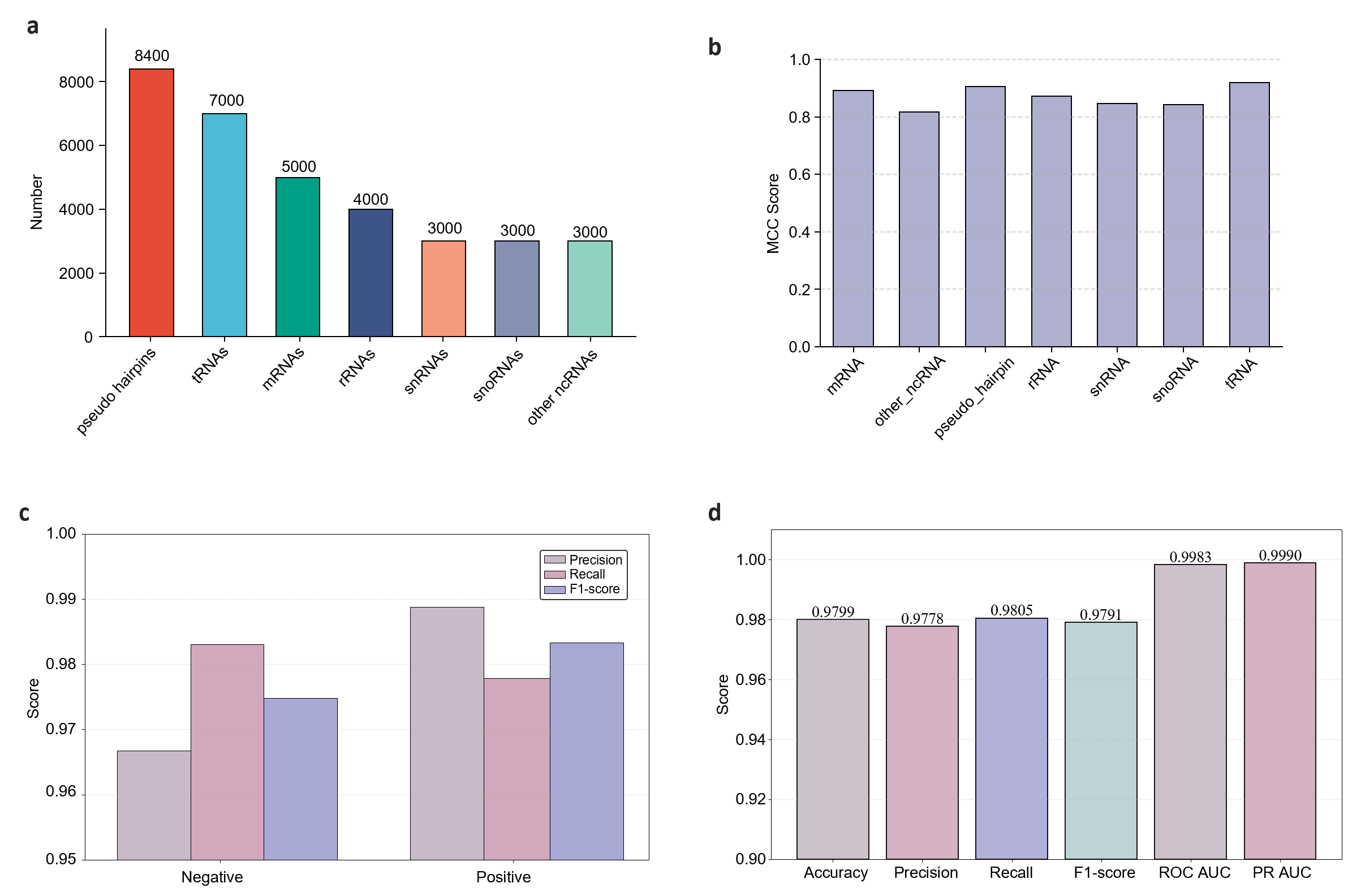

**Supplementary Figure 1 | Evaluating Binary Classification of miRNA Precursors Against Other RNAs.** **a,** Category distribution of negative sample in the binary classification dataset. **b,** Matthews correlation coefficient (MCC) for mixed-subtype evaluation, in which all positive validation samples were assessed together with one negative subtype at a time. Higher MCC indicates stronger overall agreement between predictions and true labels under each subtype-specific binary setting. **c,** Class-specific validation performance for the negative class and positive class, shown in terms of precision, recall and F1-score. **d,** Overall validation performance summarized by six metrics, including Accuracy, macro-averaged Precision, macro-averaged Recall, macro-averaged F1-score, ROC AUC and PR AUC.

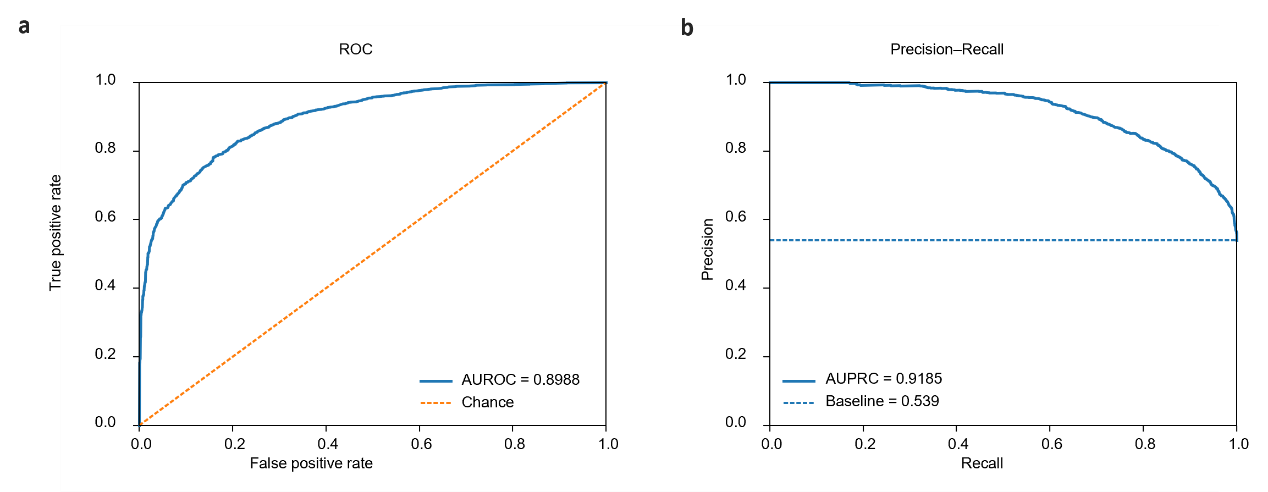

**Supplementary Figure 2 | Performance of the binary classifier of mature miRNA on the validation set.** **a,** Receiver operating characteristic (ROC) curve for predicting whether the annotated guide mature miRNA product originates from the 5′ armor 3’ arm. The dashed diagonal indicates chance performance. Area under the ROC curve (AUROC) is shown in the legend. **b,** Precision–recall (PR) curve for the same task. The dashed horizontal line indicates the baseline precision equal to the positive-class prevalence in the validation set (pos_rate ≈ 0.539). Area under the PR curve (AUPRC) is shown in the legend.

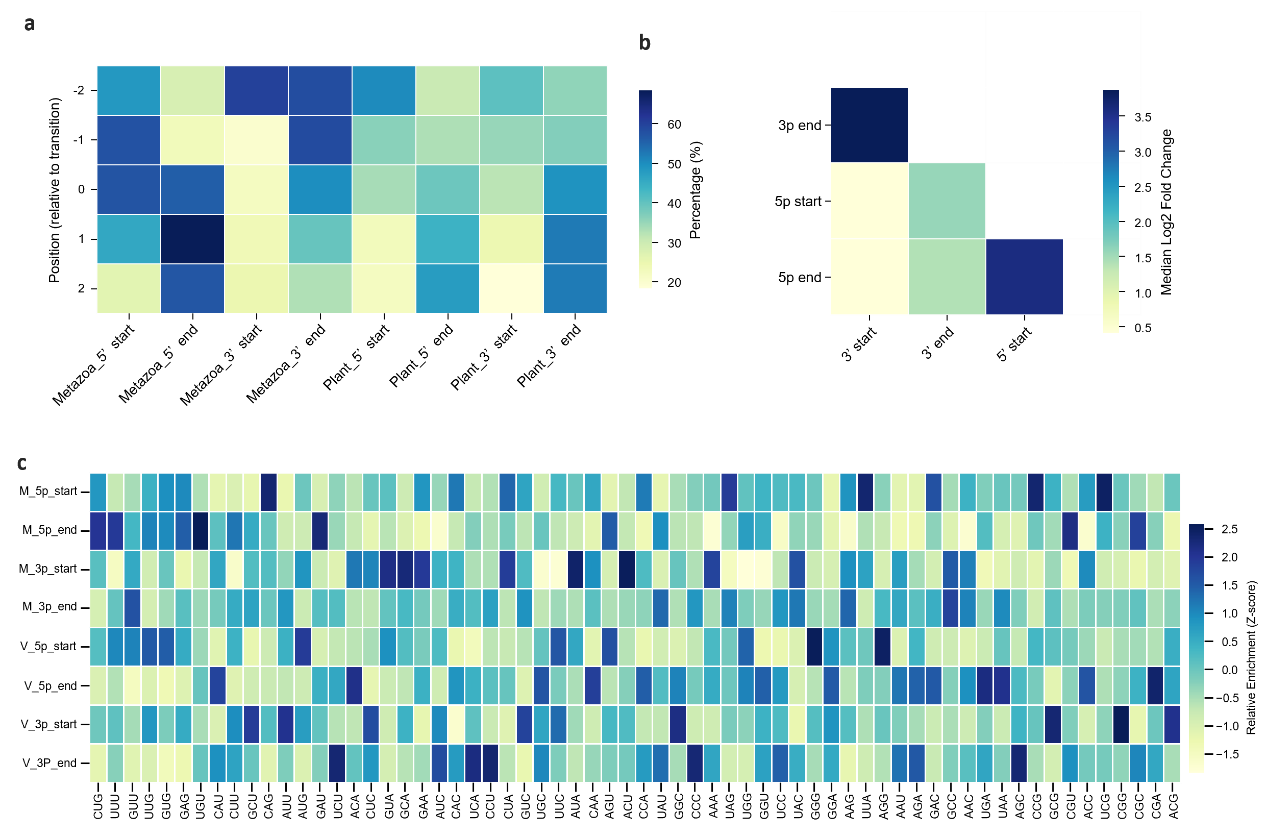

**Supplementary Figure 3 | Boundary-proximal sequence–structure patterns.** **a,** Frequency of local sequence–structure transitions around mature-miRNA duplex boundaries across clades. Heatmap showing the frequency of transition positions within a ±2 nt window around each of the four duplex boundaries (5p start, 5p end, 3p start and 3p end) in Metazoa and Viridiplantae. Columns denote the eight boundary–clade groups, and rows denote positions relative to the annotated boundary coordinate. A position was classified as a transition position when its local secondary-structure state differed from at least one immediate neighbor, corresponding to a local paired-to-unpaired or unpaired-to-paired change in the predicted precursor structure. Cell values indicate the percentage of valid precursor instances in which the corresponding relative position satisfied this criterion. This analysis resolves the positional landscape of local structural switching around duplex boundaries and highlights boundary-specific asymmetry in transition enrichment. **b,** Pairwise coupling strength between mature-miRNA duplex boundaries inferred from attention wiring. Lower-triangle heatmap summarizing pairwise attention coupling between the four duplex boundaries (3p start, 3p end, 5p start and 5p end). For each precursor, high-weight attention links extracted from the shared-encoder attention graph were classified according to whether they connected the neighborhoods of two annotated boundary types, and enrichment was quantified relative to a permutation-based null expectation. Each cell reports the median log2 fold change across sampled precursors for the corresponding boundary pair, with warmer colors indicating stronger-than-expected coupling. The lower triangle is shown to avoid redundancy. This analysis reveals how the model jointly organizes boundary-related information at the level of pairwise interactions and highlights preferential coupling between specific boundary pairs. **c,** Boundary-specific local sequence contexts around mature-miRNA duplex boundaries. Heatmap showing the distribution of local sequence motifs in the immediate vicinity of the four mature-miRNA duplex boundaries across Metazoa and Viridiplantae. For each boundary instance, a 3-nt window centered on the annotated boundary position (±1 nt) was extracted from the precursor sequence, and the most frequent motifs were ranked globally after pooling all boundary–clade groups. This visualization highlights that the four duplex boundaries are associated with distinct local sequence biases rather than a single shared boundary-proximal sequence pattern.

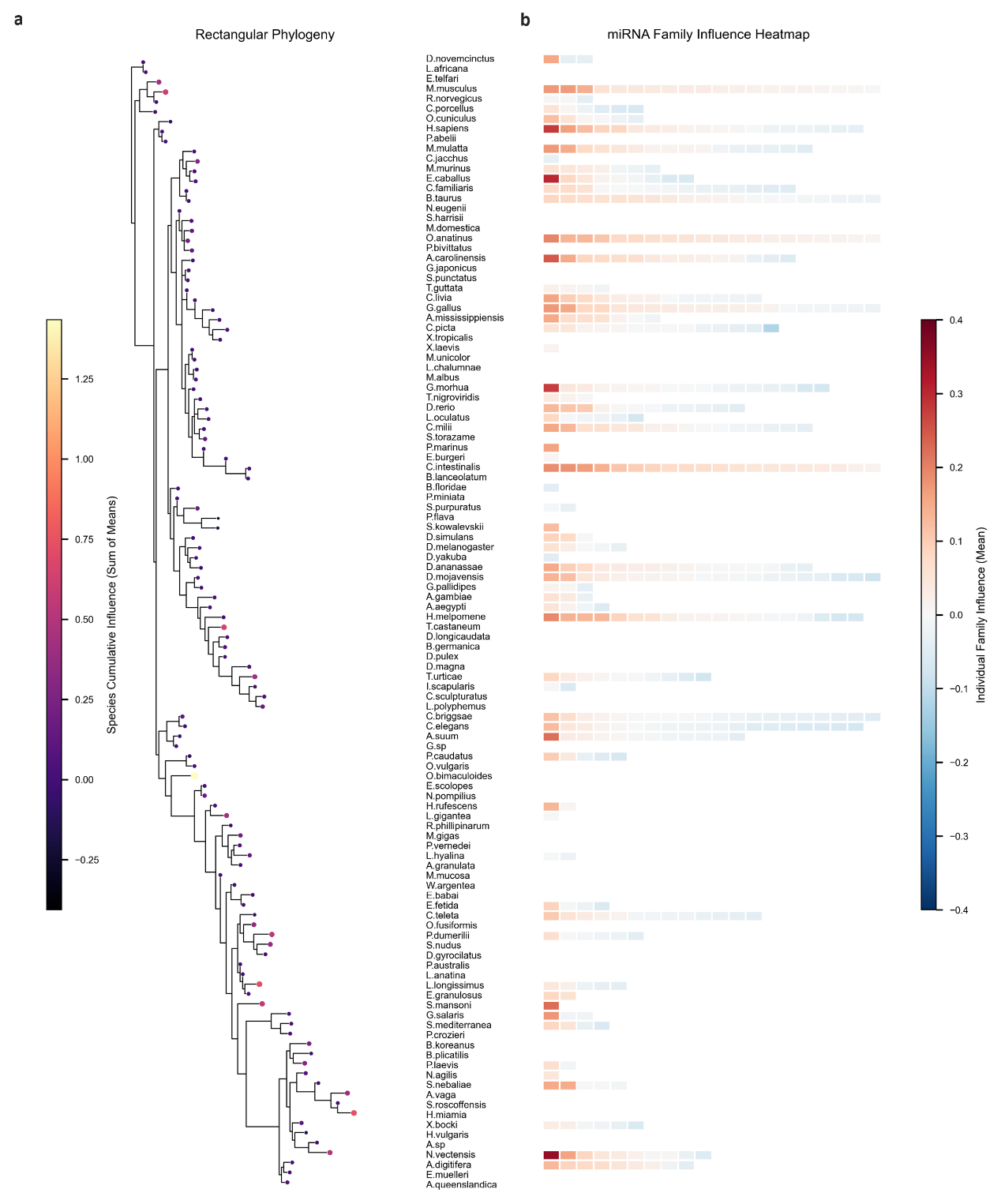

**Supplementary Figure 4 | Phylogenetic divergence and functional emergence of miRNA families.** **a,** Phylogenetic reconstruction of MirGeneDB3 species. The central phylogeny illustrates the evolutionary relationships among 114 metazoan species, with branch lengths representing substitutions per site (square-root transformed for visual clarity). Terminal nodes are marked with circular indicators, where the size and color intensity represent the cumulative net influence (sum of mean *influence*) of the miRNA families associated with each specific lineage. Critically, each node in the tree corresponds to a specific evolutionary stage where novel miRNA families emerged; the data mapped to these positions reflect the functional influence of families that were first derived at that point in the phylogeny. **b,** Comparative landscape of miRNA family influence. The peripheral heatmap displays the individual influence scores of the top 20 miRNA families for each species, aligned in a one-to-one correspondence with the terminal leaves of the phylogeny. The color gradient indicates the mean influence score, with red representing positive and blue representing negative functional influence. This integrated visualization allows for the direct correlation between the evolutionary origin of a miRNA family and its subsequent functional impact across diverse biological lineages.

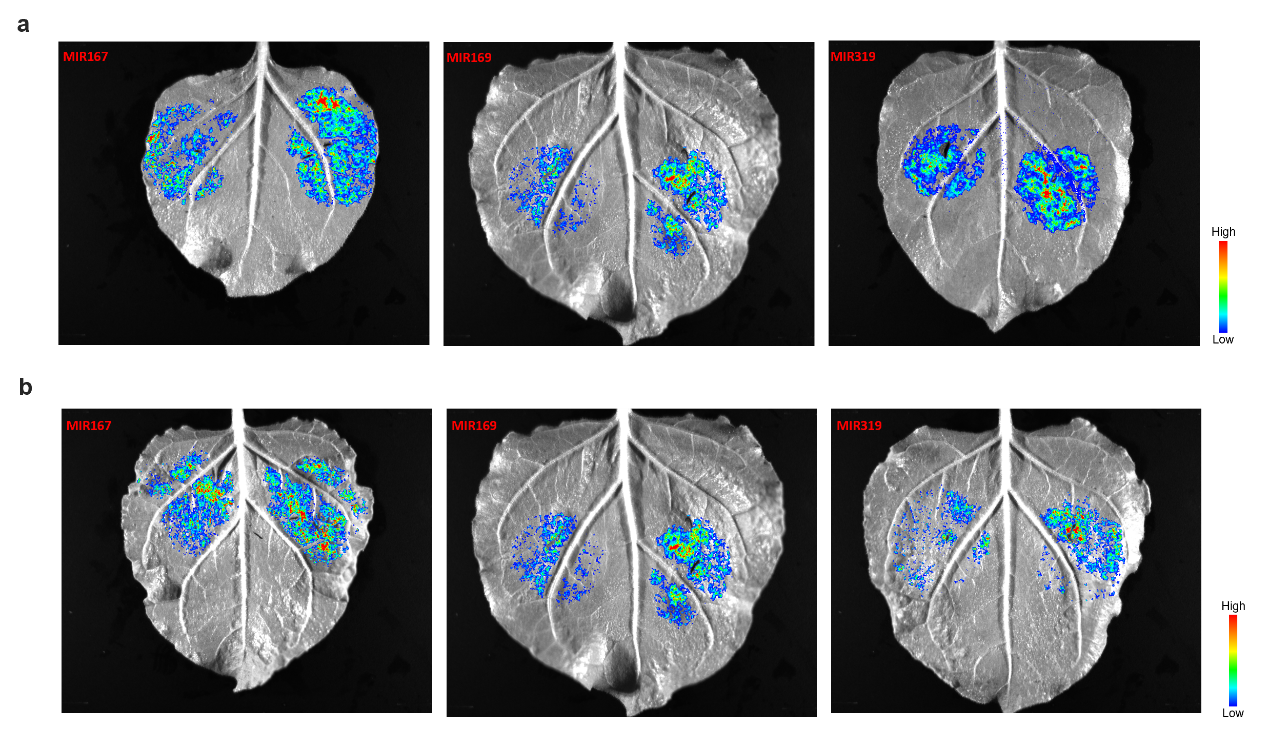

**Extended Data Fig. 5 | Luciferase reporter validation of LUC-targeting amiRNA constructs.** **a,** Representative paired luciferase (LUC) reporter assay images for amiRNA constructs targeting the LUC reporter. Three constructs are shown: MIR167, MIR169 and MIR319 scaffolds. In each leaf, the LUC reporter and the indicated amiRNA construct were co-infiltrated on one side, whereas the opposite side received the LUC reporter alone as a within-leaf control. Pseudocolor signals indicate relative LUC activity, with reduced signal in the LUC + amiRNA region indicating reporter repression. **b,** Independent replicate bioluminescence images for the same LUC-targeting amiRNA constructs shown in **a**.
